# The evolution of a Na^+^-sensitive *Vibrio cholerae* mutant unmasks the moonlighting aminopeptidase PepA as a regulator of *nhaB* Na^+^/H^+^ antiporter gene expression

**DOI:** 10.64898/2026.03.13.711655

**Authors:** Sebastian Herdan, Katharina Kohm, Robert Warneke, Florian Roth, Nicolas Görge, Trung Duc Hoang, Emina Schunke, Claudia C. Häse, Juri Rappsilber, Günter Fritz, Fabian M. Commichau, Johannes Gibhardt, Julia Steuber

**Author notes:** The authors should be regarded as joint first authors (Sebastian Herdan, Katharina Kohm).

## Abstract

The pathogenic bacterium *Vibrio cholerae* is native to seawater and can therefore be cultivated in nutrient media with increased salt concentration. To maintain osmotic balance, *V. cholerae* uses Na^+^/H^+^ antiporters and an Na^+^-translocating NADH:quinone oxidoreductase (Na^+^-NQR). The exact contribution of the various Na^+^/H^+^ antiporters to maintain Na^+^ and H^+^ homeostasis in *V. cholerae* is unclear. However, genetic studies indicate the major Na^+^/H^+^ antiporter NhaA and the Na^+^-NQR are required by the bacteria for Na^+^ resistance at alkaline conditions. Here we show that the growth defect of a *V. cholerae nhaA nqr* mutant at increased Na^+^ concentrations and alkaline pH is relieved by the rapid acquisition of suppressor mutations. The suppressor mutants could be assigned to two classes. We identified (i) mutations in the promoter of the *nhaB* gene encoding a Na^+^/H^+^ antiporter and (ii) mutations that either affect the expression level of the *pepA* gene or DNA binding activity of the encoded multifunctional aminopeptidase PepA. The characterization of the *P_nhaB_* and *P_pepA_* promoters of the suppressor mutants, membrane potential measurements and comparative proteome analyses identified PepA as a novel factor controlling Na^+^ homeostasis in *V. cholerae*.

**IMPORTANCE:** The emergence and spread of multi-resistant bacteria are a major problem for humans and animals. Therefore, the identification of novel targets for the development of novel antibiotics to combat pathogenic bacteria is extremely important. Since sodium ion homeostasis is an essential process in many pathogenic bacteria, especially those living in marine environments, it represents an interesting target for antibiotics. The perturbation of sodium ion homeostasis does indeed impair the fitness of the human pathogenic bacterium *Vibrio cholerae* but can be restored by the rapid evolution of the bacteria. This suggests that inhibiting multiple targets with different antibiotics might be more effective in preventing the development of resistant bacteria.

## INTRODUCTION

To maintain osmotic balance, all living cells must be capable of extruding sodium ions (Na^+^) from the cytoplasm to the extracellular space. Moreover, bacteria like alkaliphilic *Bacilli* and marine *Vibrio* use Na^+^-driven flagellar motor complexes for locomotion in their aquatic habitats [1]. To prevent the accumulation of Na^+^, these bacteria rely on transport systems for the export of the cation. The extrusion of Na^+^ can be achieved with the help of Na^+^/H^+^ antiporters, which are secondary transporters that couple the export of Na^+^ to the import of H^+^ [2–8]. For instance, in *Escherichia coli*, the export of Na^+^ is driven by at least two distinct Na^+^/H^+^ antiporters, NhaA and NhaB, of which the former is required for the maintenance of the cytoplasmatic pH at alkaline pH [9–13]. The expression of the *nhaA* gene is positively controlled by the LysR-type DNA-binding transcription factor NhaR that responds to Na^+^ [14–17]. NhaA couples the export of 2 Na^+^ to the import of 3 H^+^ and therefore is crucial for acidification of the cytoplasm under alkaline conditions [18]. Moreover, some bacteria can establish an electrochemical sodium gradient using primary Na^+^ pumps, such as the Na^+^-translocating NADH:quinone oxidoreductase (Na^+^-NQR) [19], or the Na^+^-pumping oxaloacetate decarboxylase [20]. The Na^+^-NQR is a six-subunit, respiratory complex that couples the oxidation of NADH and reduction of ubiquinone to the translocation of Na^+^ across the inner bacterial membrane [21]. In *Vibrio cholerae*, the causative agent of Cholera disease, the Na^+^-NQR operates together with several Na^+^/H^+^ antiporters [6,22] to ensure survival in very different habitats, ranging from alkaline marine environment with high salinity to acidic compartments in the human gastrointestinal tract [23,24]. Moreover, virulence and pathogenicity of *V. cholerae* are intimately linked to the activity of the Na^+^-NQR [23,25], marking the protein complex and other respiratory enzymes of pathogenic bacteria as putative drug targets [26,27].

In addition to the Na^+^-NQR, five antiporter systems have been identified in *V. cholerae* that contribute to Na^+^ and H^+^ homeostasis [19]. For instance, the Na^+^/H^+^ antiporter NhaA in *V. cholerae* [28], is the homolog of NhaA from *E. coli* [29]. Like in *E. coli*, the expression of the *nhaA* gene depends on the Na^+^-responsive transcriptional activator NhaR [30]. Moreover, NhaB, of which also a homolog exists in *E. coli*, is a Na^+^/H^+^ antiporter that confers resistance to Na^+^ and Li^+^ when overproduced in an *E. coli nhaA nhaB* mutant [6,31]. The K^+^(Na^+^)/H^+^ antiporter NhaP1 also seems to be required for pH homeostasis in *V. cholerae* [32]. Furthermore, the multi-subunit Na^+^/H^+^ antiporter Mrp allowed the *E. coli nhaA nhaB* mutant to grow at toxic Na^+^ and Li^+^ concentrations as well as with natural bile salts [33]. While the exact contribution of the Na^+^/H^+^ antiporters NhaB, NhaD, NhaP1 and Mrp in maintaining Na^+^ and H^+^ homeostasis in *V. cholerae* is yet unclear, NhaA can be considered as the major Na^+^/H^+^ antiporter that, together with the Na^+^-NQR, was shown to be required by the bacteria for Na^+^ resistance at alkaline conditions [6].

In the present study, we confirmed that the *nhaA* and *nqr* genes encoding the Na^+^/H^+^ antiporter NhaA and Na^+^-NQR, respectively, become synthetically lethal in *V. cholerae* at high Na^+^ concentrations and pH. Serendipitously, we found that the *V. cholerae nhaA nqr* mutant is genomically highly unstable and rapidly forms suppressor mutants in which Na^+^/H^+^ homeostasis is restored. The characterization of the suppressor mutants revealed that restoration of Na^+^/H^+^ homeostasis can be achieved by the strong overexpression of the *nhaB* Na^+^/H^+^ antiporter gene. The suppressor screen also identified the multifunctional PepA protein as a novel repressor of the *nhaB* gene because loss-of-function mutations in *pepA* results in NhaB overproduction and Na^+^ resistance. As an aminopeptidase, PepA is involved in amino acid metabolism, and the protein’s DNA binding activity controls plasmid segregation and gene expression. The role of PepA in maintaining Na^+^/H^+^ homeostasis in *V. cholerae* is discussed.

## RESULTS

### Genomic stability of a *V. cholerae nqr nhaA* mutant at high salinity and alkaline pH

Previously it was observed that the mutational inactivation of the *nhaA*, *nhaB* and *nhaD* Na^+^/H^+^ antiporter genes in *V. cholerae* did not affect exponential growth of the bacteria [6]. Moreover, a *V. cholerae nqr* mutant lacking the NQR complex did not show defects in Na^+^-pumping-related phenotypes [34]. This suggests that other secondary Na^+^ pumps can compensate for the lack of NQR. Indeed, the selective inhibition of the NQR by 2-*n*-nonyl-4-hydroxyquinoline *N*-oxide uncovered that the NhaA Na^+^/H^+^ antiporter becomes essential for Na^+^ resistance at alkaline conditions [6]. Since the NQR is an interesting target for novel antibiotics, we assessed the genomic stability of a *V. cholerae* strain lacking the major Na^+^/H^+^ antiporter and the primary pump. For this purpose, we cultivated the wild type, the *nhaA* and *nqr* single mutants, and the *nhaA nqr* double mutant in standard LB medium (pH 6.2, 171 mM NaCl) or in LB with Tris-buffered pH (pH 7.8) without additional NaCl (residual Na^+^ concentration, 13 mM). As shown in **Figures 1A** and **1B**, the strains lacking the NQR produced slightly less biomass during stationary growth phase. This indicates some contribution to Na^+^ extrusion by the primary Na^+^ pump under these conditions. In contrast, *nhaA nqr* mutant showed a severe growth defect in alkaline LB medium containing 150 mM NaCl (**Figure 1C**). As expected, the growth defect of the *nhaA nqr* mutant can be relieved by plasmid-based overexpression of the *nqr* operon (**Figure 1D**). Thus, in perfect agreement with earlier studies, the *nhaA* and *nqr* genes are synthetically lethal in *V. cholerae* at high salinity and alkaline pH [6,35]. However, after 17 h of growth a transition to almost wild type growth rate was observed for the *nhaA nqr* mutant (**Figure 1C**). This indicates that this mutant is genomically unstable and theresulting selective pressure leads to rapid emergence of suppressor mutations that confer Na^+^ resistance.

**Figure 1.**
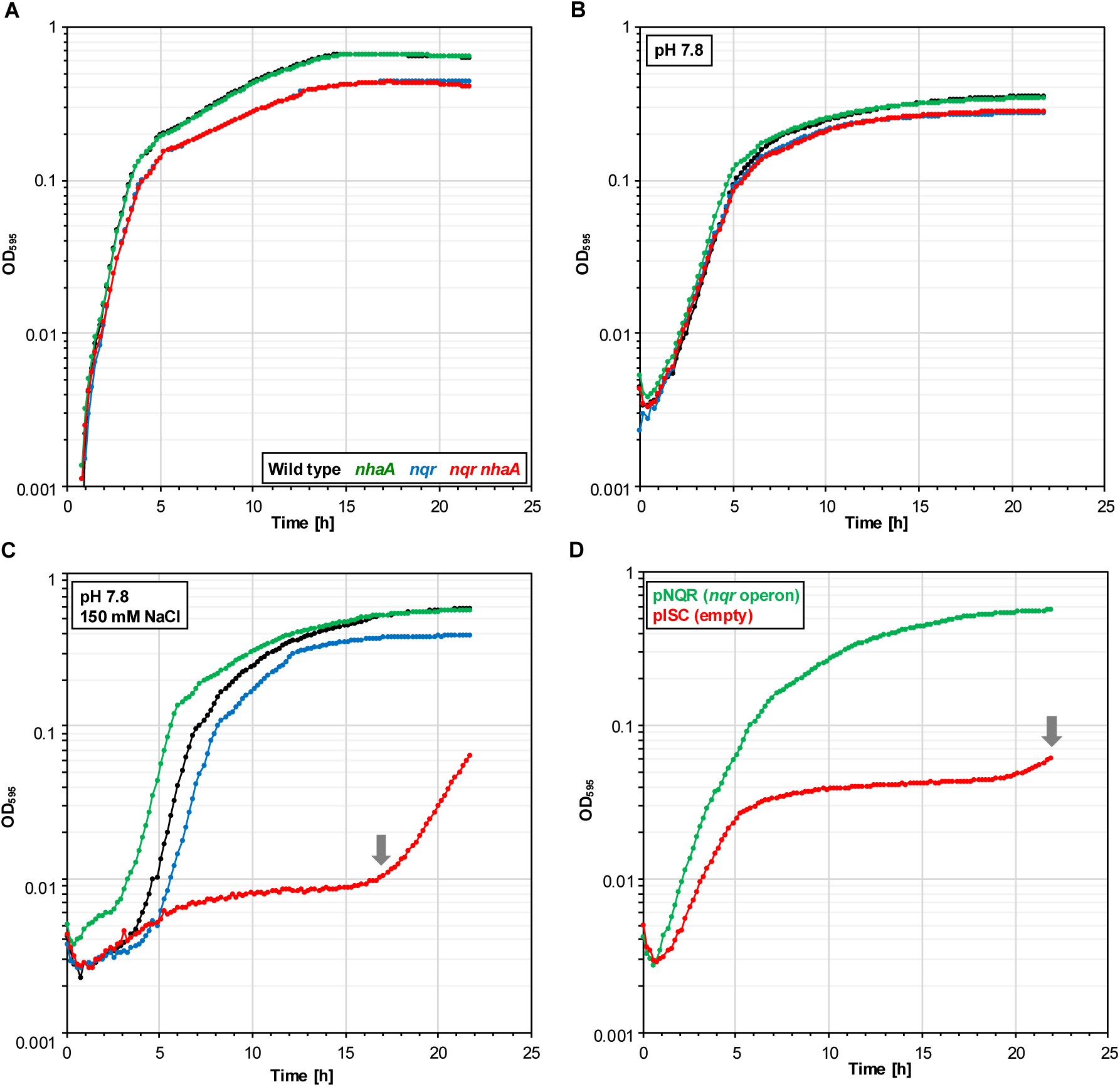
Influence of salinity and pH on growth of *V. cholerae* mutants with impaired Na^+^ export. Growth of the wild type and the *nhaA*, *nqr* and *nqr nhaA* mutants in LB medium (pH 6.2, 171 mM NaCl) (**A**), in LB_Tris_ (pH 7.8, 13 mM Na^+^) (**B**), and in LB_Tris_ (pH 7.8) in the presence of 150 mM NaCl (**C**). Growth of the *nqr nhaA* mutant carrying the empty plasmid pISC or the plasmid pNQR for the overexpression of the NQR in LB_Tris_ medium at pH 7.8 in the presence of 150 mM NaCl (**D**). The cultivation experiments were carried out in a multi-well plate reader at 30 °C. The grey arrows indicate the emergence of suppressor mutants.

### Characterization of *V. cholerae nqr nhaA* suppressor mutants

We repeatedly observed that the *V. cholerae nqr nhaA* mutant when grown in LB medium at high salinity and pH 7.8 resumed growth (**Figures 1C** and **1D**; data not shown). As described above, this indicates the formation of Na^+^-resistant suppressor mutants that probably overproduce a Na^+^ extrusion system. To understand the underlying molecular mechanism of Na^+^ homeostasis restoration in the background of the *V. cholerae nqr nhaA* mutant, we isolated suppressor mutants. For this purpose, we grew the mutant overnight in standard LB medium (pH 6.1, 171 mM NaCl), washed the cells twice in standard LB and propagated the bacteria on selective LB plates (pH 7.8, 150 mM NaCl). After incubation for 5 days at 30°C, approximately 250 suppressor mutants appeared on the plates on which 10^7^ cells had been distributed (**Figure 2A**). Next, we isolated 20 independent suppressors (S1 – S20) (Table 1) and verified the genetic stability of the mutants by passaging of the bacteria on standard LB plates and subsequent selection at high salinity and alkaline pH. The cultivation of a selected set of mutants (S3, S4 and S7) in selective LB medium confirmed their Na^+^ resistance. As shown in **Figure 2A**, even though the mutants produced less biomass, their growth rates were comparable to that of the wild type strain. The determination of the membrane potential (**Figure 2B**) revealed that all mutants were less impaired in formation of membrane potential than the parental *V. cholerae nqr nhaA* strain. Furthermore, potentials of the mutants were at least as high as the ones of the *V. cholerae nqr* strain, thus resembling rescue from the *nhaA* deletion in the parental strain. To conclude, the *V. cholerae nqr nhaA* mutant is genetically unstable and quickly adapts to high salinity and alkaline pH through the accumulation of beneficial mutations.

**Figure 2.**
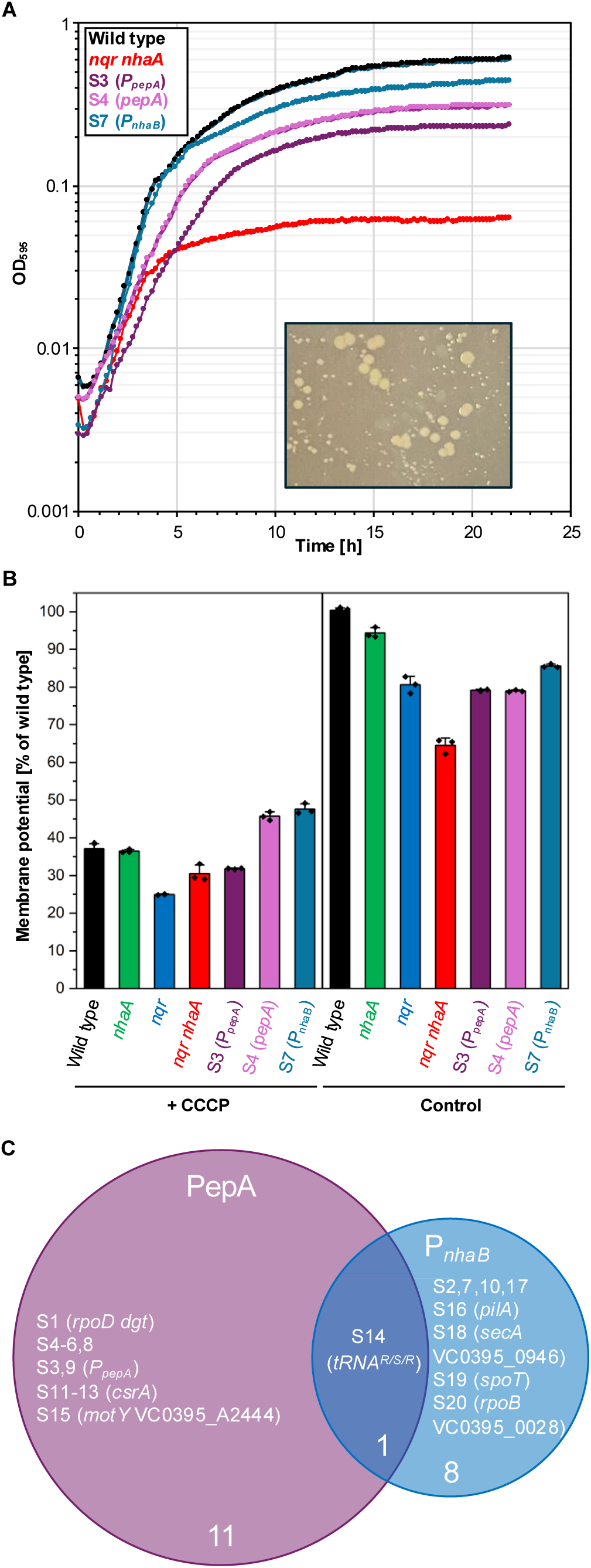
Characterisation of selected suppressor mutants and identified genomic alterations relieving the growth defect of the *nqr nhaA* mutant. (**A**) Growth of the wild type, the *nhaA*, *nqr* mutant, and the suppressor mutants S3, S4 and S7 in LB medium at pH 7.8 in the presence of 150 mM NaCl. The cultivation experiments were carried out in a multi-well plate reader at 30 °C. The Figure inlay shows the suppressor mutants that emerged during growth under selection. (**B**) Determination of the membrane potential in the wild type, the *nhaA*, mutant, the *nqr* mutant, the *nhaA*, *nqr* mutant, and the suppressor mutants S3, S4 and S7. (**C**) Venn diagram illustrating the overlap of the genomic regions mutated in the *nqr nhaA* strain during growth under selection (see also Table 1).

**Table 1.**
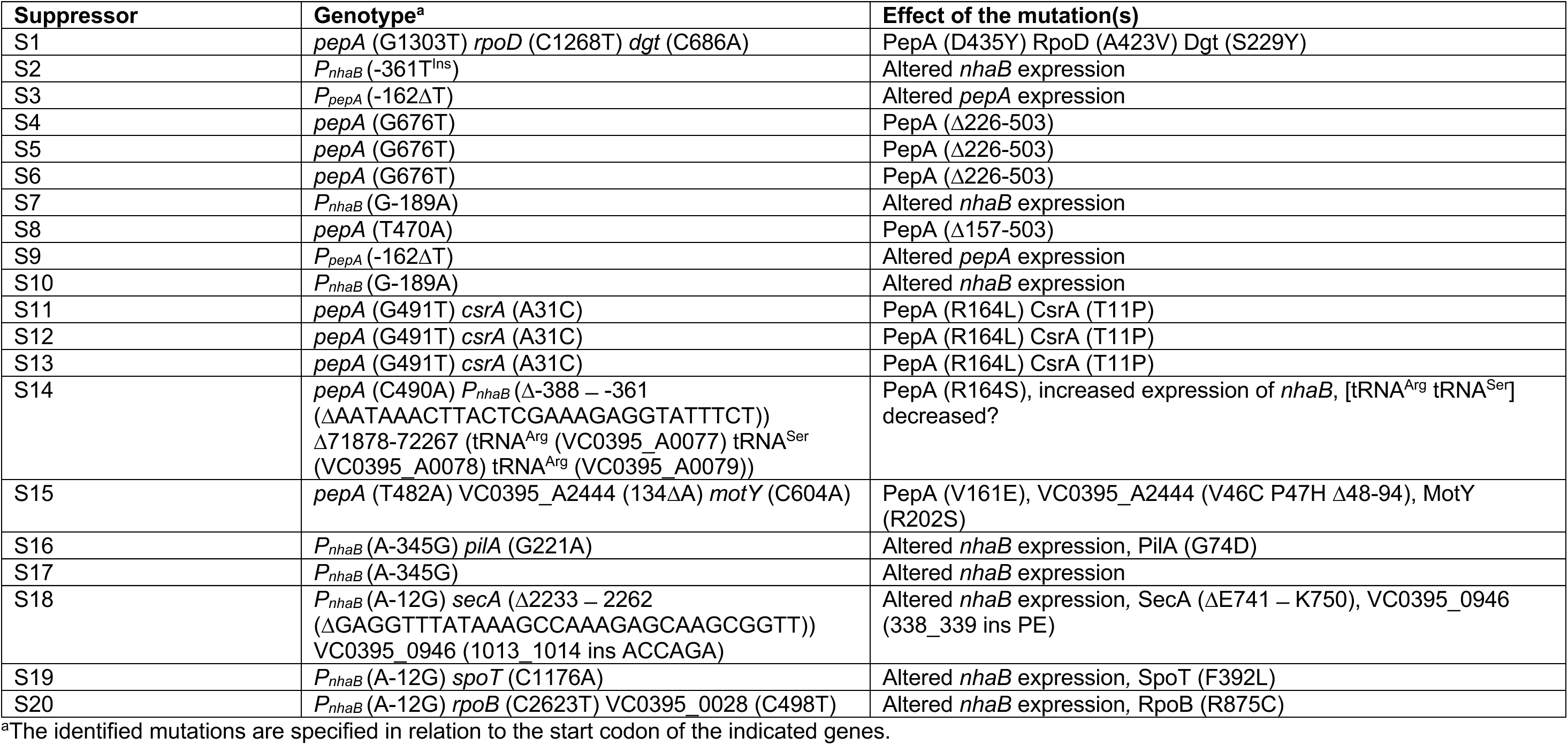
Isolated *V. cholerae nqr nhaA* suppressor mutants.

### Identification of the mutations in the *nqr nhaA* suppressors

Genome sequencing of the 20 isolated suppressors allowed us to assign the mutants to two different classes. Nine mutants had acquired a mutation in a genomic region that likely contains the promoters of the *nhaB* and *fadR* genes (**Figure 2C** and **3A**). The *fadR* gene encodes a DNA-binding transcription factor that is involved in fatty acid degradation and biosynthesis [36,37]. Since NhaB is a Na^+^/H^+^ antiporter, the mutations in the *fadR-nhaB* intergenic region likely increase the activity of the *P_nhaB_* promoter, which result in overproduction of NhaB and thus Na^+^ resistance of the *V. cholerae nqr nhaA* suppressors. Interestingly, three *P_nhaB_* promoter variants emerged independently of each other twice (**Figure 3A**) (Table 1). The remaining 12 suppressor mutants all had acquired mutations that either affect the expression levels of the *lptF* and *pepA* genes, or the coding sequence of the *pepA* gene (**Figure 2C, 4A**) (Table 1). The suppressor S14 with a mutation in the *pepA* gene also had deleted a 27 bp-long sequence in the *fadR-nhaB* intergenic region. Recently, chromatin immuno precipitation sequencing (ChIP-Seq) experiments revealed that PepA is a global transcription repressor in *V. cholerae* [38]. Interestingly, the *fadR-nhaB* and *lptF-pepA* intergenic regions were also identified as PepA binding sites (**Figure 3A, Figure S2**). This suggests that the mutations in the *fadR-nhaB* and *lptF-pepA* intergenic regions may abolish the PepA-dependent repression of the *P_nhaB_* promoter and affect negative autoregulation and thus the cellular concentration of the repressor.

**Figure 3.**
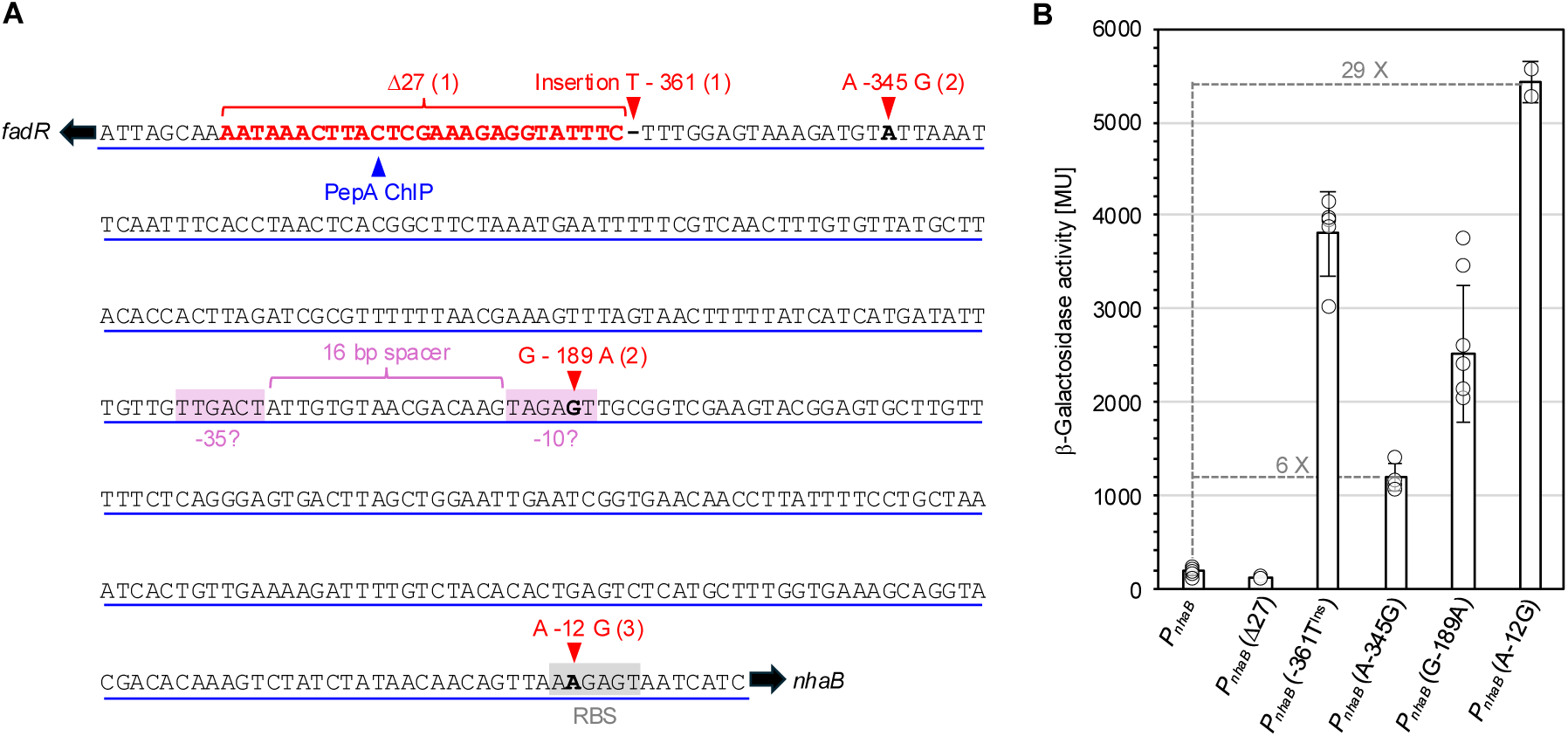
Characterization of the *P_nhaB_* promoter variants of the *nqr nhaA* suppressors. (**A**) *fadR-nhaB* intergenic region located on chromosome 1 of *V. cholerae.* The identified mutations are indicated in red and are specified in relation to the start codon of the *nhaB* gene. The PepA binding site in the *P_nhaB_* promoter is underlined and the region that was most enriched in the ChIP-Seq experiment (14-fold) is indicated by a blue arrow [38]. RBS, ribosome-binding site. A possible promoter created or improved by the mutation A −189 A mutation is highlighted in pink. (**B**) β-Galactosidase assay to assess the effect of the mutations in the *fadR-nhaB* intergenic region on the activity of the *P_nhaB_* promoter in *E. coli* XL1-Blue. The cells were grown in LB medium at 37 °C and harvested at an OD_600_ between 0.5 and 0.8. β-Galactosidase activities are given as units per milligram protein. Experiments were carried out at least three-fold. Mean values are indicated by columns.

As with the first class of mutants, two *pepA* mutations were also identified multiple times (Table 1). Moreover, five *pepA* mutants had acquired additional mutations in genes whose coding products are involved in post-transcriptional regulation (CsrA), Na^+^-driven motility (MotY), cell surface structure (PilA), stringent response (SpoT), information processing (Dgt, RpoB, RpoD), and protein secretion (SecA), translation (tRNA^Arg^ tRNA^Ser^), and unknown processes (VC0395_0946, VC0395_A2444) [39–42]. However, the fact that in six mutants of the second large group of suppressors only the *pepA* gene or the *lptF-pepA* intergenic region were altered suggests that the encoded protein is involved in the control of Na^+^ extrusion in *V. cholerae*. In summary, the link between PepA and Na^+^ resistance in the *nqr nhaA* suppressors appears less obvious at first glance. However, it is often the case that different mutations produce similiar phenotypes in suppressor screens [43–45].

### *nhaB* overexpression restores the growth defect of the *nqr nhaA* mutant

To assess whether the mutations in the *fadR-nhaB* intergenic region of the *V. cholerae nqr nhaA* suppressors affect the activity of the *P_nhaB_* promoter, we constructed translational *nhaB-lacZ* fusions and introduced the plasmids into *E. coli* XL1-Blue. Unfortunately, a *lacZ*-based reporter system to assess the activity of the *P_nhaB_* promoter could not be used in our *V. cholerae* strain because it already contains a *lacZ* gene in the *toxT* locus (Table 1). However, since *E. coli* and *V. cholerae* are both *γ*-proteobacteria, we assumed that the wild type *P_nhaB_* promoter and the mutant derivatives must also be active in *E. coli*. It has indeed been shown that a *V. cholerae* DNA-binding regulator can replace the functional homolog in *E. coli* [30]. Next, the strains were grown in LB medium, and the cells were harvested during mid-exponential growth (OD_600_ 0.5 – 0.8). Except for the promoter with the 27 bp-long deletion (derived from suppressor S14), all promoters with single nucleotide exchanges were 6 – 29-fold more active than the wild type *P_nhaB_* promoter (**Figure 3B**). In suppressor S14, the mutation in *pepA* likely causes de-repression of the *P_nhaB_* promoter in *V. cholerae* (see below). To conclude, it is reasonable to assume that, due to the mutations in the *fadR-nhaB* intergenic region, the *V. cholerae nqr nhaA* suppressors overproduce Na^+^/H^+^ antiporter NhaB, which leads to Na^+^ resistance through increased export of the cation.

### A potential link between DNA-binding activity of PepA and *nhaB* gene expression

The remaining 12 *nqr nhaA* suppressor mutants all had acquired mutations that either affect the expression level or the coding sequence of the *pepA* gene (**Figure 2C**) (Table 1). In two suppressors (S3 and S9), the same nucleotide was deleted in the *lptF-pepA* intergenic region (**Figure 4A**). While *lptF* codes for an essential component of a lipopolysaccharide transport system, the *pepA* aminopeptidase gene is not essential in *V. cholerae* [38,46]. Therefore, it is unlikely that the mutation in the suppressors S3 and S9 would affect *lptF* expression. To assess the wild type and mutant *P_lptF_* and *P_pepA_* promoters, we constructed translational *lptF*-and *pepA-lacZ* fusions and introduced the plasmids into *E. coli* XL1-Blue. Next, we propagated the transformants on LB plates containing X-Gal. The *E. coli* cells carrying the wild type and mutated *lptF-lacZ* fusions both formed dark blue colonies. The β-galactosidase assay confirmed that the mutation in the *lptF-pepA* intergenic region does not affect the activity of the *P_lptF_* promoter (**Figure 4B, 4C**). In contrast, cells with the mutated *pepA-lacZ* fusion formed white colonies on the plates compared to the cells with the wild type-promoter (**Figure 4B**). This made it more difficult to identify positive clones using blue-white screening. The β-galactosidase test revealed that the mutated *pepA-lacZ* fusion was about 17-fold less active (**Figure 4C**). In summary, the mutation in the *P_pepA_* promoter of the suppressors S3 and S9 likely leads to a reduction of the cellular PepA concentration. Using the *E. coli pepA* wild type and *pepA::tet* strains, we tested whether *E. coli* PepA has an influence on the *nhaB-lacZ* fusion. However, this was not the case (data not shown).

**Figure 4.**
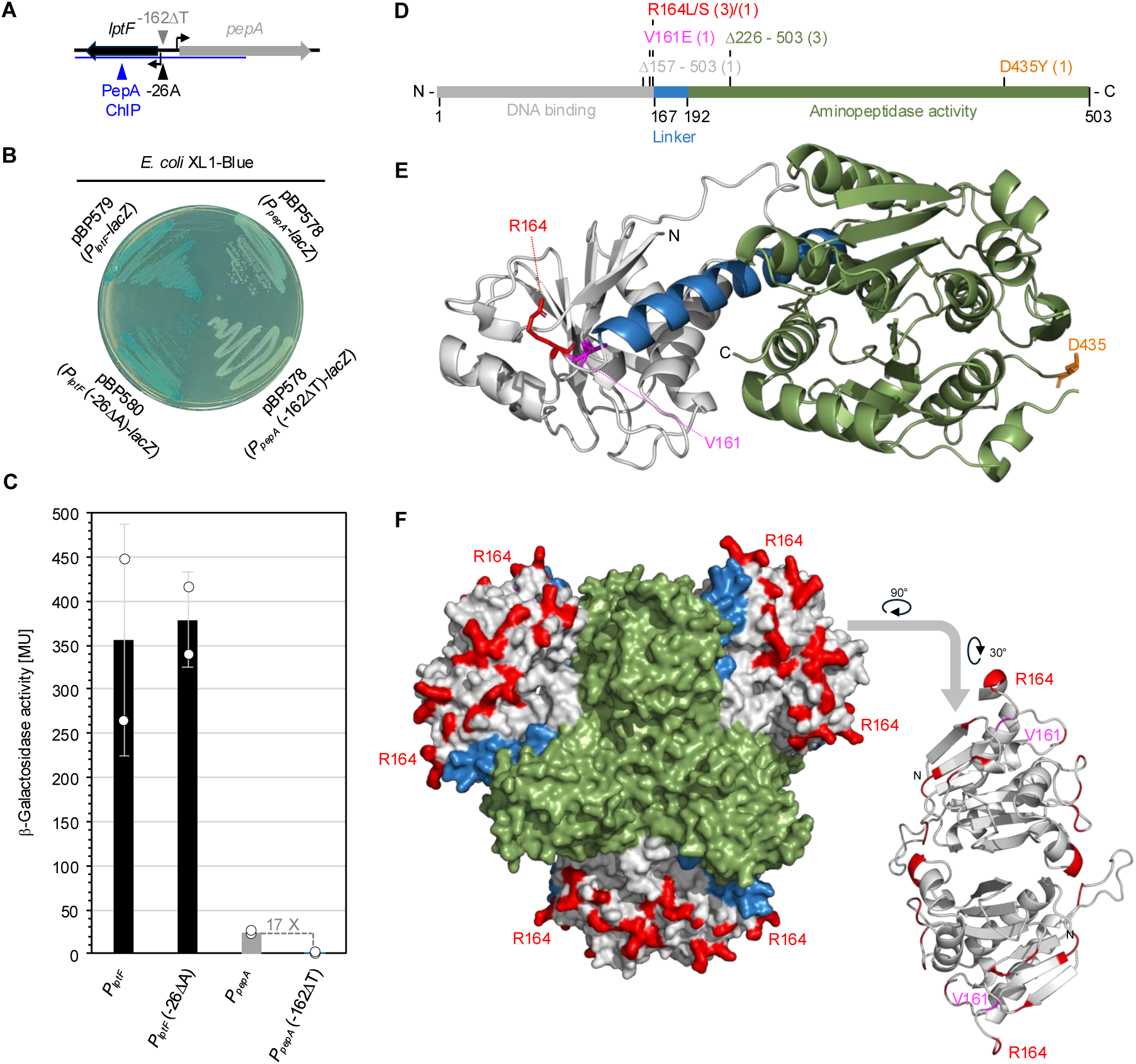
Characterization of the *P_pepA_* promoter variant and PepA variants in the *nqr nhaA* suppressors. (**A**) *lptF-pepA* intergenic region located on chromosome 1 of *V. cholerae.* The identified mutation is specified in relation to the start codons of the genes. The PepA binding site in the *lptF-pepA* intergenic region is underlined and the region that was most enriched in the ChIP-Seq experiment is indicated by a blue arrow [38] (**B**) Qualitative test to visualize the activities of the *P_pepA_* and *P_lptF_* promoters in *E. coli* XL1-Blue that was propagated on LB medium containing the chromogenic substrate X-Gal. The plates were incubated for 24 h at 37 °C. (**C**) β-Galactosidase assay to assess the effect of the mutation in the *lptF-pepA* intergenic region on the activity of the *P_pepA_* and *P_lptF_* promoters in *E. coli* XL1-Blue. The cells were grown in LB medium at 37 °C and harvested at an OD_600_ between 0.5 and 0.8. β-Galactosidase activities are given as units per milligram protein. Experiments were carried out at least three-fold. Mean values are indicated by columns. (**D**) Amino acid replacements in PepA. The numbers in parentheses indicate how often the PepA variants were identified. (**C**) Localization of the amino acid exchanges in a full-length PepA model based on the structure of the *E. coli* homolog (PDBid: 1GYT). (**F**) Molecular surface of the *V. cholerae* PepA hexamer. The amino acid residues that were previously identified to be important for DNA-binding activity and XerCD-dependent recombination are labelled in red [48].

The remaining 10 suppressors carry mutations in *pepA* that likely affect the DNA-binding activity of the encoded protein (**Figure 2C, 4D**) (Table 1). For instance, four suppressors (S4, S5, S6 and S8) had acquired a stop codon that would truncate PepA by 228 or 347 amino acids (**Figure 4B**). An extensive characterization of PepA mutant variants from *E. coli* revealed that the N-terminal domain (residues 1 – 166) is involved in DNA binding, whereas the C-terminal domain (residues 193 – 503) that is connected by an α-helix (residues 167 – 192) is required for aminopeptidase activity (**Figure 4D, 4E**) [47,48]. Thus, the truncation in S4 – S6 and S8 probably leads to a general defect in PepA folding or oligomerization and thus to the loss of DNA-binding and aminopeptidase activities. The remaining six suppressors contain *pepA* alleles that code for PepA variants with single amino acid exchanges, among them four with replacements of arginine 164 (**Figure 4D, 4E**). The suppressor S15 synthesizes a PepA variant in which a residue located near arginine 164 in PepA was replaced. The arginine 164 was previously identified as crucial for the DNA-binding activity of *E. coli* PepA [48]. Therefore, the exchange of arginine 164 and neighboring residues likely affects the DNA-binding activity of *V. cholerae* PepA. The suppressor S1 synthesizes a PepA variant in which aspartate 435 is replaced by tyrosine (**Figure 4D, 4E**). Even though the aminopeptidase domain is altered in this PepA variant, changes in the C-terminal domain can lead to both enzyme functions being impaired [48]. As described above, PepA is a global transcription factor that has many chromosomal binding sites, including the *P_nhaB_* and *P_pepA_* promoters (**Figure 3A, 4A**) [38]. Moreover, suppressor S14 had deleted a part of the PepA binding site in the *P_nhaB_* promoter and synthesizes a PepA variant in which the arginine 164 is replaced by leucine (**Figure 3A, 4D, 4E**). Thus, the synthesis of the PepA R164L variant and the deletion of the PepA binding site in the *P_nhaB_* promoter likely have a synergistic effect on *nhaB* expression in *V. cholerae*. In summary, the suppressor analysis suggests that PepA is involved in Na^+^ homeostasis by controlling the expression of the *nhaB* Na^+^/H^+^ antiporter gene.

### Identification of PepA in the *nqr nhaA* suppressors and PepA-regulated genes

To evaluate how the reduced PepA synthesis, the synthesis of a truncated PepA variant, or the increased *P_nhaB_* promoter activity affect the global physiology of the *nqr nhaA* suppressors, we performed comparative proteome analyses. For this purpose, we cultivated the parental strain and the suppressors S3 (*P_pepA_*), S5 (*pepA*) and S7 (*P_nhaB_*) in LB medium under non-selective conditions. As a control, we also compared the proteomes of an *E. coli* strain and its isogenic *pepA* mutant. It is known that PepA represses the *carAB* operon that encodes the CarAB carbamoylphosphate synthetase complex, which is involved in pyrimidine biosynthesis [47,49–51]. Therefore, we expected that there would be less of CarA and CarB in *E. coli* and possibly also in the *V. cholerae* suppressors S3 and S5. Indeed, both proteins were much less abundant in the *E. coli pepA* mutant and in the two *nqr nhaA* suppressors (**Figure 5A**). Interestingly, in the *E. coli pepA* mutant, the PyrIB enzyme complex was less abundant as compared to the parental strain. PyrIB is also involved in pyrimidine biosynthesis and catalyzes the reaction that follows CarAB. In the *V. cholerae* mutants S3 (*P_pepA_* (−162ΔT)) and S4 (*pepA* (G676T)), as expected, PepA abundance was strongly reduced. In contrast, suppressor S7 (*P_nhaB_* (G-189A); **Figure 5B**) showed no changes in PepA abundance. Furthermore, the comparative proteome analyses suggest different metabolic strategies of the suppressor mutants to counteract Na^+^ stress (**Figure 5B**, supplemental material Table S1). For example, in suppressor S7 carrying a mutation in the *nhaB* promotor, which results in increased *nhaB* expression (**Figure 3B**), there is a more than two-fold increase in levels of citrate lyase subunits CitF and CitC and oxaloacetate decarboxylase subunits OadA-2 and OadG, which are part of the *cit* operon allowing citrate fermentation [52]. Moreover, the malate synthase AceB is highly overexpressed, likely further facilitating oxaloacetate biosynthesis from isocitrate via the glyoxylate cycle. This will increase overall Na^+^ export capacity with help of the OadA-2 sodium pump [20] and might decrease carbon flux into the TCA cycle. In contrast, in suppressors S3 (*P_pepA_* (−162ΔT)) and S4 (*pepA* (G676T)), in which either expression of the *pepA* promoter is impaired (**Figure 4C**) or truncated PepA is expressed, carbon flux through the TCA cycle might be increased. Both the citrate lyase subunits (CitF in S3 and S4 and in S4 also CitD) and OadA-2 (S3 and S4) are less abundant in these mutants, while at the same time there is a moderate increase in citrate synthase subunit GltA in S4. Furthermore, in S4 the fumarase FumC is moderately less abundant, while the succinate-ubiquinone oxidoreductase complex SdhAB is moderately more abundant. Interestingly, in line with increased abundance of TCA cycle enzymes, enzymes involved in amino acid biosynthetic pathways, especially for glutamate, tryptophan, glutamine, histidine and others are also more abundant. Moreover, moderately increased levels of the ATP synthase are observed. This suggests that an increase in membrane potential due to respiratory activity in suppressor S3 and especially S4 facilitates Na^+^ export by the remaining Na^+^/H^+^ antiporters. This is probably accompanied by changes in the membrane structure in suppressor S4, as indicated by alterations in the protein levels of cell-envelope related proteins such as NlpC, CpxP, MinCD (Supplemental material Table S1). However, the shapes of the *V cholerae* reference strain, and of *V. cholerae nqr nhaA* and its suppressor strains, did not differ significantly (**Supplemental material Figure 1**). Changes in abundance of the Na^+^/H^+^ antiporter NhaB itself could not be identified. This is not surprising because NhaB is an integral membrane protein with low recovery. To conclude, the proteome analyses revealed that the mutation in the *P_pepA_* promoter (S3) and in *pepA* (S4), reduce the cellular amount of PepA. Moreover, PepA also regulates *carAB* expression in *V. cholerae* and changes in the proteomes indicated different adaptation strategies of the *pepA* and *nhaB* suppressor mutants.

**Figure 5.**
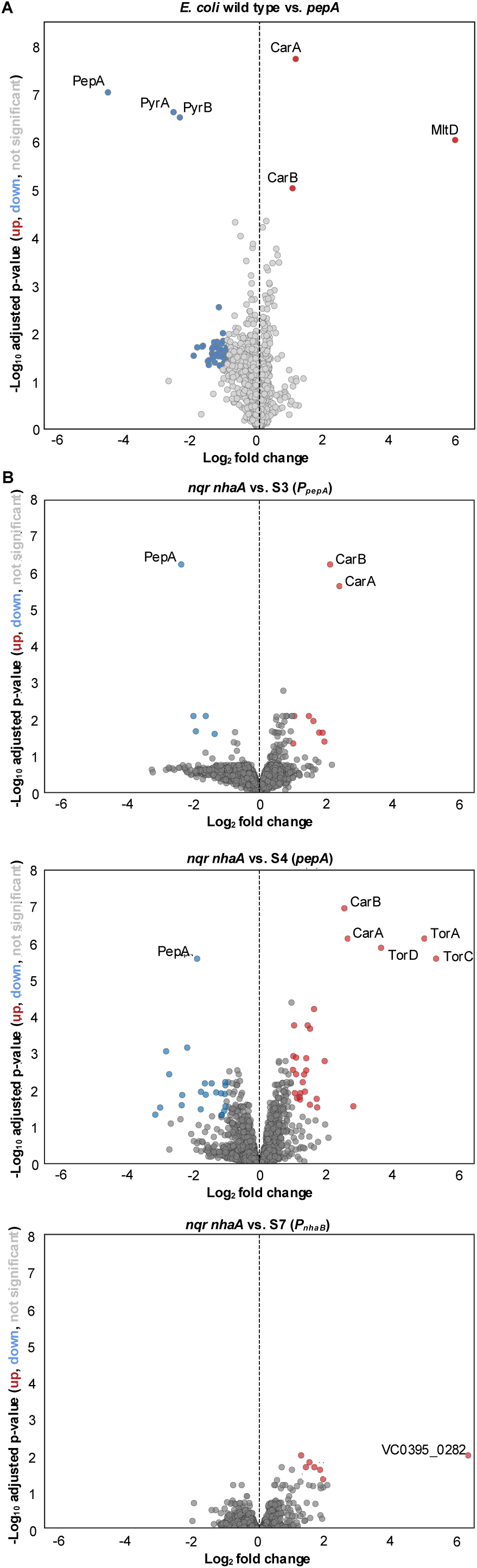
Comparative proteome profiling and chromosomal PepA binding sites. Proteome comparisons of the *E. coli* wild type (DS941) and *pepA* mutant (SDC1) strains (**A**) and of the *V. cholerae nqr nhaA* mutant and suppressors S3, S4 and S7 (**B**).

## Discussion

In the present study, we found that the *V. cholerae nqr nhaA* mutant with impaired Na^+^/H^+^ homeostasis is genomically unstable during growth in the presence of elevated Na^+^ concentrations at alkaline pH. Similarly, like the *V. cholerae nqr nhaA* mutant, an *E. coli nhaA nhaB* mutant lacking the two specific Na^+^/H^+^ antiporters rapidly forms suppressor mutants at high salinity [53]. The characterization of one of these suppressor mutants revealed that the Na^+^ export capacity was partially restored and the growth rate was like that of the wild type. Although the genomic alteration allowing the growth of this mutant at elevated Na^+^ concentrations was not identified, it was suggested that under some conditions *E. coli* can synthesize a primary Na^+^ pump when the Na^+^/H^+^ antiporters NhaA and NhaB are not present [53]. It is tempting to speculate that the *E. coli nhaA nhaB* mutant was relieved by a mutation that would affect the synthesis of the other known Na^+^/H^+^ antiporters like ChaA, MdfA and MdtM [54–56]. However, in contrast to the Na^+^-sensitive *E. coli nhaA nhaB* mutant, in *V. cholerae*, the overproduction of the Na^+^/H^+^ antiporter NhaB can be achieved either by overexpression of the *nhaB* gene or by mutations that affect the synthesis of PepA, a multifunctional protein that has been extensively studied in *E. coli*. PepA was first identified as an aminopeptidase that allows *E. coli* to utilize peptides as nutrient sources [57]. Interestingly, PepA is also a component of a plasmid dimer resolution system that contributes to the segregational and inheritable stability of natural plasmids like ColE1 [58]. PepA, together with the DNA-binding regulator or arginine biosynthesis ArgR, serve as accessory proteins, ensuring that the XerCD recombinase-dependent plasmid dimer to monomer conversion occurs exclusively unidirectional [50,59–66]. Recently, it has been observed that PepA might even serve as an RNA-binding chaperone in *E. coli* [67]. Moreover, PepA and ArgR are required for stable maintenance of the P1 prophage and the repression of the *carAB* operon in *E. coli* [47,49–51,68]. Thus, PepA can be assigned to the class of trigger enzymes, a group of moonlighting proteins that are active in metabolism and in controlling gene expression [69,70]. The regulation of gene expression by trigger enzymes can be achieved by direct binding to DNA or RNA, by modulating the activity of a DNA-binding transcription factor or by covalent modification of a transcription factor [69,70]. In many cases, the regulatory activity of a trigger enzymes is controlled by the presence or absence of a substrate or cofactor. For instance, the aconitase catalyses the reversible conversion of citrate to isocitrate in the tricarboxylic acid cycle and requires an iron-sulphur cluster for activity [71]. Under conditions of iron limitation, the aconitase binds to iron-responsive elements in the mRNAs of genes involved in iron homeostasis [72]. However, in contrast to the aconitase, PepA does not require any pyrimidine cofactor to bind to DNA *in vitro* [49]. Moreover, the structural characterisation and the identification of PepA variants with single and multiple amino acid exchanges revealed that the peptidase activity of the protein is not required for its DNA-binding and XerCD site-specific recombination activity [47–49,73–76].

Recently, in *E. coli* and *V. cholerae* it has been shown that the available carbon source affects the cellular localization of PepA and its DNA-binding activity [38]. In the presence of glucose, a carbon source that allows rapid growth, PepA is sequestered to the membrane by a direct interaction with the global transcription regulator CAP [38]. During nutrient-limited growth (e.g., growth with acetate as the carbon source), PepA is released from the membrane and becomes active as a DNA-binding transcription factor. It has been hypothesized that the release of PepA from the membrane tailors a stress response to a nutrient-limited environment [38]. ChIP-seq analyses identified 293 binding sites for PepA on the two chromosomes of *V. cholerae* O1 biovar El Tor str. N16961, among them the *P_nhaB_* and *P_pepA_* promoter regions (**Figure 3A, 4A** and **supplemental material Figure 2**) [Gibson et al., 2022]. Our suppressor analysis supports the finding that PepA is a regulator of gene expression and Na^+^/H^+^ homeostasis in *V. cholerae* (**Figure 6**). The results of our comparative proteome study obtained with derivatives of *V. cholerae* O395 compare favorably with the findings of Gibson *et al*. 2022 [38], who identified PepA DNA binding sites in *V. cholerae* N16961. In many cases, PepA binding revealed by ChlP-seq occurred upstream of genes, which are also affected in our suppressor strains, as identified by proteomics (Table S1). In other cases, changes in protein abundance are likely needed to redirect the flux of metabolites, e.g. towards the nitrogen metabolism to prevent bottlenecks or imbalances due to high expression of e.g. CarAB. Some proteins with higher abundance that may directly regulated by PepA are, for example, the glutamate synthase GltB and PurDH enzyme complex (moderately increased) that are involved in purine biosynthesis. Their expression likely needs to be coordinated with CarAB to prevent metabolic imbalances. PepA also appears to be involved in the control of toxin gene expression in *V. cholerae* O395 [77]. Thus, PepA links carbon source availability to virulence in *V. cholerae*, a phenomenon that has been observed in many other pathogenic bacteria [78].

**Figure 6.**
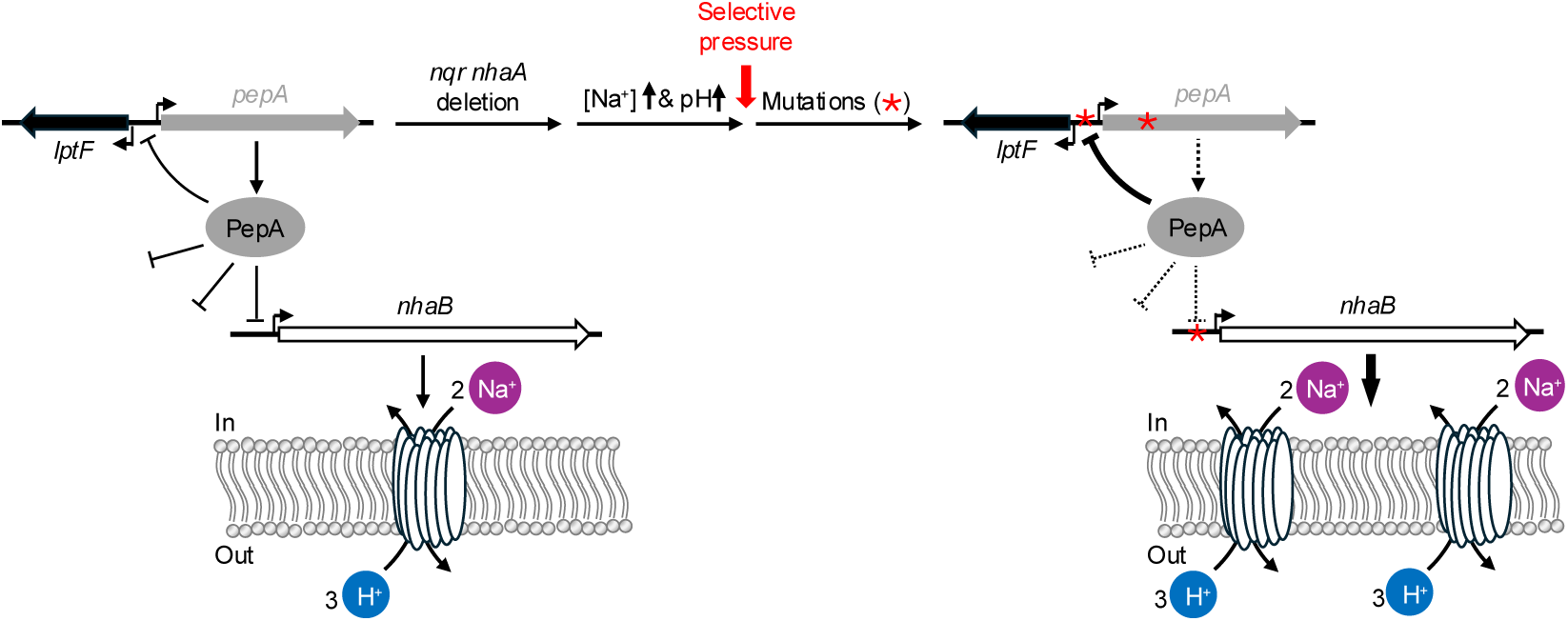
Model for the regulation of the *P_nhaB_* promoter by PepA. The cultivation of the *nhaA nqr* mutant under selection (high salinity and alkaline pH) causes the accumulation of mutations that affect *pepA* expression, the cellular amount of PepA, and *nhaB* expression.

As for *V. cholerae,* the PepA binding landscape in *E. coli* was globally mapped by ChIP-seq [38,79]. In contrast to *V. cholerae*, *E. coli* PepA belongs to the group of transcription factors and only binds to two promotors, thereby controlling its own synthesis and the expression of the *carAB* operon [49,80]. The influence of PepA on the cellular concentrations of the PyrIB complex and the membrane-bound lytic murein transglycosylase D (MtlD) is novel and deserves further investigation (**Figure 5A**). In *Vibrio anguillarum*, MtlD has been associate with virulence of the fish pathogen [81]. To conclude, PepA is a fascinating protein, and further investigations will be necessary to decipher the significance of this multitasking protein for the global bacterial physiology.

## MATERIALS AND METHODS

### Bacterial strains and chemicals

Bacteria and oligonucleotides (purchased from Sigma-Aldrich, Munich, Germany) used in this study are listed in Table 2. Chemicals and media were purchased from Sigma-Aldrich (Munich, Germany), Carl Roth (Karlsruhe, Germany) and Becton Dickinson (Heidelberg, Germany). Bacterial chromosomal DNA was isolated using the peqGOLD bacterial DNA kit (Peqlab, Erlangen, Germany). PCR products were purified using the PCR purification kit (Qiagen, Germany; New England Biolabs). Phusion DNA polymerase was purchased from Thermo Scientific (Germany) and used according to the manufacturer’s instructions.

**Table 2.**
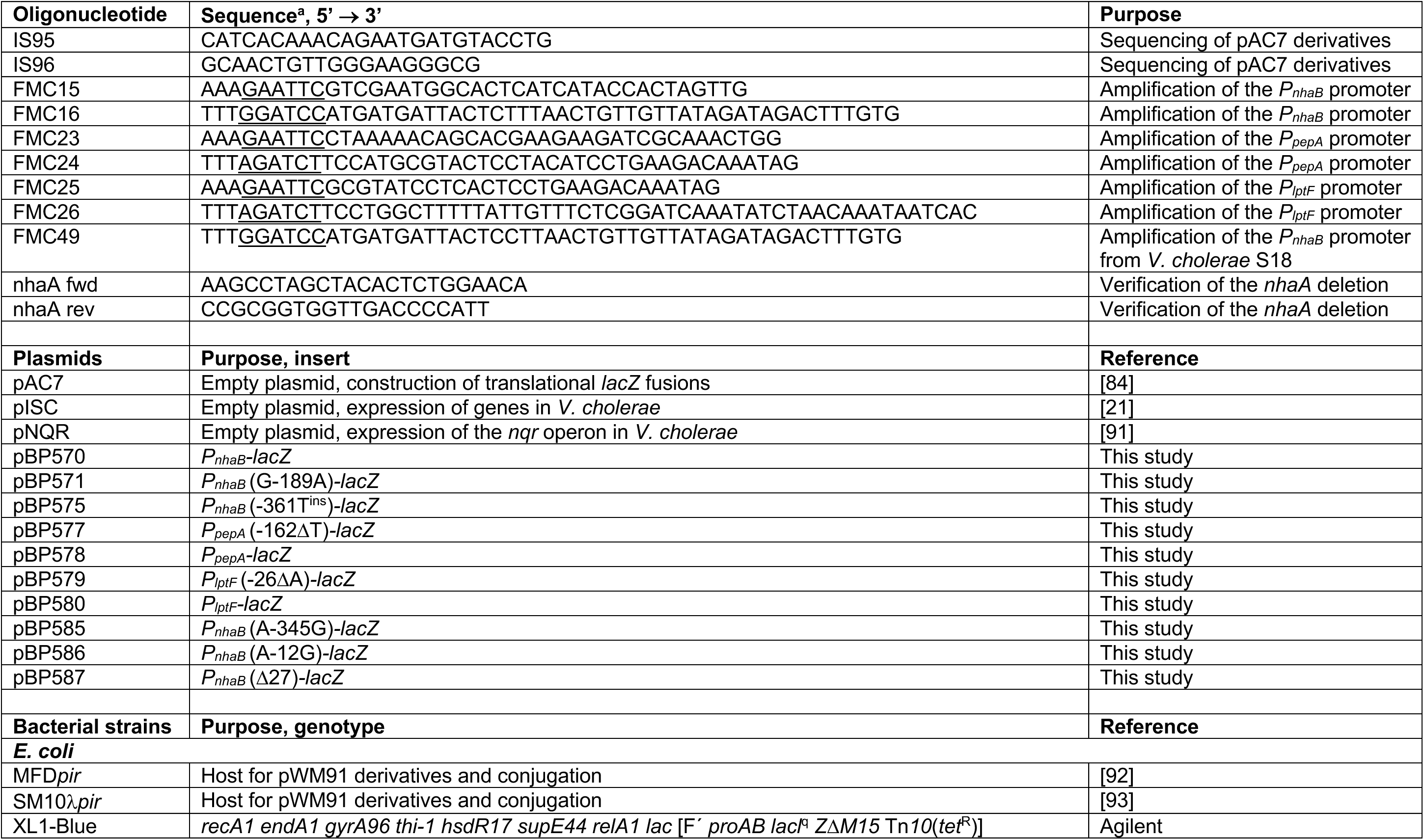
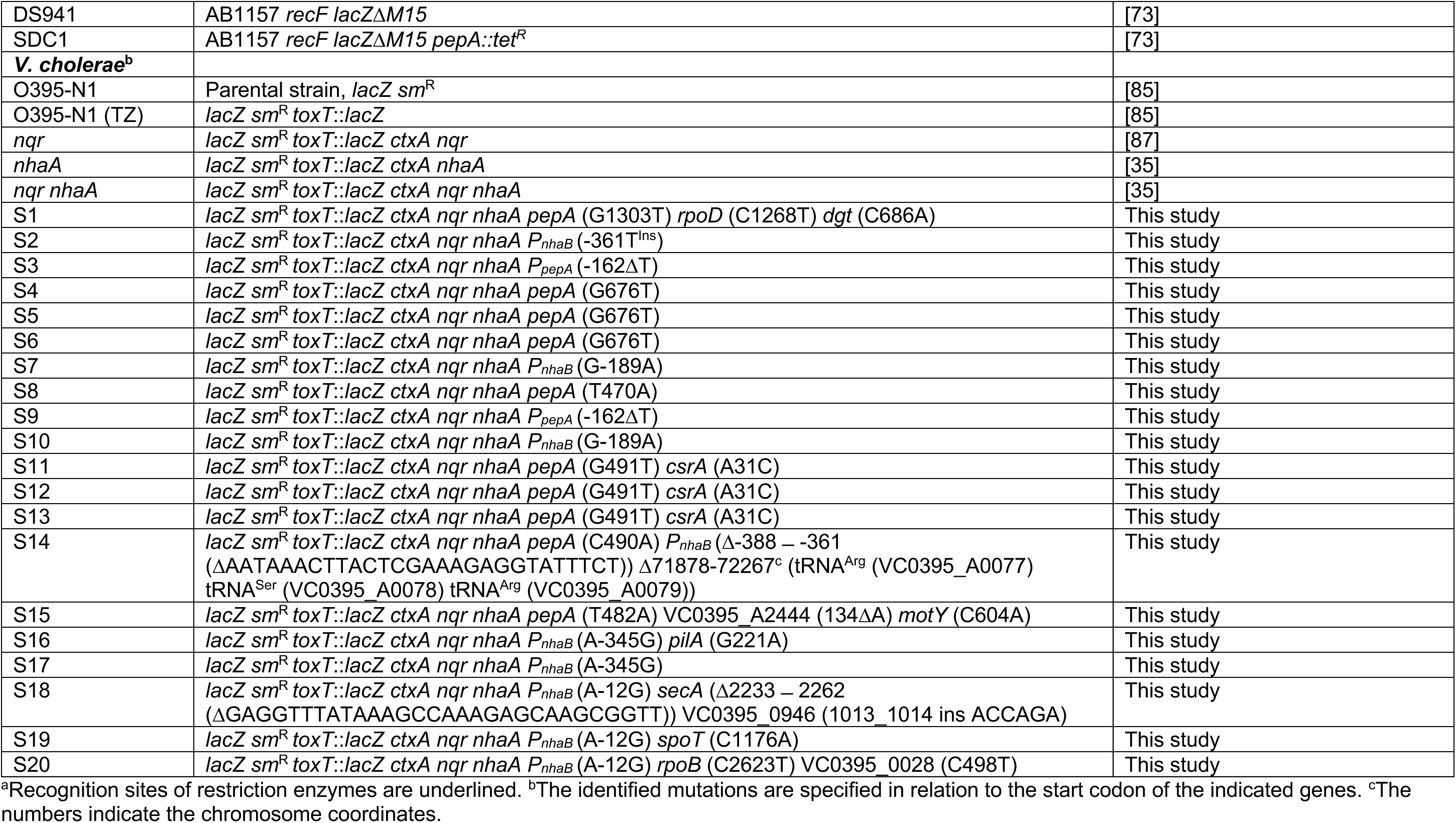
Oligonucleotides, plasmids and bacterial strains.

### Cultivation of *E. coli* and *V. cholerae*

*E. coli* was grown in lysogeny broth (LB) at 37°C [82,83]. Agar plates were prepared with 15 g agar/l (Roth, Germany). *E. coli* cells that were transformed with the plasmid pAC7 and its derivatives were selected and LB plates supplemented with 100 µg/ml ampicillin and 80 µg/ml X-Gal. *V. cholerae* was grown in LB, LB_sucrose_, LB_Tris_, and LB_Tris_-NaCl medium. LB medium contains 1 % tryptone (w/v), 0.5 % yeast extract (w/v), 171 mM NaCl and has a pH of 6.2. LB_sucrose_ contains 1 % tryptone (w/v), 0.5 % yeast extract (w/v) and 10 % sucrose (w/v)]. LB_Tris_ medium was used to monitor the effect of Na^+^ on the growth and contains 100 mM Tris. The pH was adjusted to 7.6 with HCl at room temperature, which corresponds to pH 7.8 at 30 °C. The Na^+^ concentration of this medium was determined to be 13 mM using atomic absorption spectroscopy. LB_Tris_-NaCl medium was supplemented with 150 mM NaCl. *V. cholerae* strains were grown at 30°C and selected with 50 µg/ml streptomycin. Growth of *V. cholerae* strains in liquid medium at 30°C was monitored in 24-well plates (Thermo Scientific, USA) in an Infinite F200 Pro plate reader (Tecan, Switzerland), and the OD_595_ was measured in 10 min intervals for 24 h. For this purpose, the *V. cholerae* strains were cultivated overnight in LB_Tris_ medium at 180 rpm and 30°C. The cultures were normalized to an OD_600_ of 0.8. Next, a 24-well-plate was prepared with 1 ml of growth medium in triplicates. After inoculation with 10 µl (1 %) of the normalized overnight culture, growth was monitored with the plate reader.

### Plasmid construction

Plasmids used and constructed in this study are listed in Table 2. Translational promoter-*lacZ* fusions for determining the activities of the *P_nhaB_*, *P_pepA_* and *P_pepA_* wild type and mutated promoters were constructed as follows. The *P_nhaB_*, *P_pepA_* and *P_lptF_* promoters were amplified by PCR using the oligonucleotide pairs FMC15/FMC16, FMC23/FMC24 and FMC25/FMC26, respectively. The primer pair FMC15/FMC49 was used to amplify the *P_nhaB_* promoter variant from suppressor S18. The *P_nhaB_* promoter fragments (615 bp) were digested with *Eco*RI and *Bam*HI and ligated with pAC7 [84] that was cut with the same enzymes. *P_pepA_* and *P_lptF_* (193 bp) promoter fragments were digested with *Eco*RI and *Bgl*II and ligated to the *Eco*RI/*Bam*HI-linearized pAC7 plasmid. All generated plasmids were verified by Sanger sequencing (Microsynth-SeqLab Sequence Laboratories, Göttingen, Germany).

### Construction of *V. cholerae nhaA* and *nqr nhaA* mutants

The nontoxic variant of the *V. cholerae* Ogawa 395 strain (O395-N1) denoted “wild type” [85,86] and the *nqr* mutant, a derivative of the strain O395-N1, lacking the *ABCDEF* operon [87] were used as parental strains (Table 2). The plasmid pWM91 *nhaA* was isolated from the *E. coli* strain SM10λ*pir* and used for transformation of the *E. coli* strain MFD*pir*. To obtain a chromosomal deletion of the *nhaA* gene in *V. cholerae*, plasmid pWM91 *nhaA* was transferred to the *V. cholerae* wild type and *nqr* mutant by conjugation using the *E. coli* donor strain MFD*pir*-pWM91 *nhaA*. The *V. cholerae nhaA* and *V. cholerae nqr nhaA* mutants were selected on LB_sucrose_ medium and deletion of the *nhaA* gene was confirmed by colony PCR with the oligonucleotide pair nhaA fwd/nhaA rev.

### Membrane potential measurements

Membrane potential of *V. cholerae* cells was measured using the BacLight bacterial membrane potential kit (Invitrogen) as described before [88]. Briefly, bacteria were grown in LB_Tris_-NaCl (30 °C, 180 rpm shaking) for 4 h, harvested (20 000 x g, 5 min, room temperature), washed in PBS (137 mM NaCl, 2.7 mM KCl, 10 mM Na_2_HPO_4_, 1.8 mM K_2_HPO_4_, pH 7.4) and adjusted to an OD_600_ of 0.25. Depolarisation control was achieved by addition of 5 µM CCCP (carbonyl cyanide 3-chlorophenylhydrazone) and incubation for 15 min in the dark at 22 °C. Cell suspensions without DiOC_2_ (3,3′-diethyloxa-carbocyanine iodide) and with DMSO served as controls. Staining was performed with 30 µM of DiOC_2_. Samples (200 µL) from three different cultures per strain were measured in black-bottomed 96-well plates (polystyrene, 4titude) in a Tecan infinite F200 Pro plate reader (excitation: 485 nm / 20 nm: emission: 535 nm / 25 nm, 635 nm / 35 nm). Background fluorescence of cell suspension without DiOC_2_ was subtracted. Fluorescence intensities were normalised to that obtained with the wild type strain.

### Genome sequencing and plotting of PepA binding sites

Genomic DNA was prepared from 2 ml overnight cultures (LB_Tris_-NaCl) using the peqGOLD Bacterial DNA Kit (VWR, USA) following the instruction manual. Purified genomic DNA was paired end sequenced (2 x 150 bp) (Azenta, Leipzig, Germany). Sequencing reads were mapped to the *V. cholerae* genome (chromosome 1, CP000626.1; chromosome 2, CP000627.1) using the Geneious (v. 2025.2.2, Docmatics) software package [89]. Genomic regions of PepA binding sites (chromatin immune precipitation (ChIP) experiments published by [38], Table S2) of *V. cholerae* O1 biovar El Tor str. N16961 (NC_002505.1 for chromosome 1 and NC_002506.1 for chromosome 2) were extracted and mapped to the genome of *V. cholerae* O395-N1 (CP000627 for chromosome 1 and CP000626 for chromosome 2) using the Geneious (v. 2025.2.2, Docmatics) software package (89 % similarity (cost matrix 93 % similarity (5.0/-9.026168) and „only transfer best match” settings) [89].

### β-Galactosidase assay

Quantitative studies of *lacZ* expression in *E. coli* were performed as follows: cells were grown at 37 °C in LB Medium. Cells were harvested at an optical density OD_600_ of 0.6 to 0.8. Specific β-galactosidase activities were determined with cell extracts obtained by lysozyme treatment as described previously [90]. One unit of β-galactosidase is defined as the amount of enzyme which produces 1 nmol of *o*-nitrophenol per min at 28 °C. The BioRad dye-binding assay was used to determine the protein concentrations.

### Electron microscopy

Bacterial cultures were grown at 30 °C in LB medium for 4 h. Cells were harvested (13 000 x g, 5 min, 4 °C) and resuspended in PBS. Samples were adjusted to an OD_600_ of 0.1 and washed twice in PBS. Cells were fixed in fixative solution (50 mM HEPES pH 7.2, 1 % p-formaldehyde, 2.5 % glutaraldehyde) for at least 2 h. After fixation, samples were gently washed three times in PBS (13 000 x g, 5 min, 4 °C) and applied on poly-L-lysine coated glass-slides. After stepwise dehydration in an ethanol series (5 min in each 30 %, 50 %, 70 %, 80 %, 90 %, 96 % EtOH), samples were critical point dried in CO_2_ (K850 Critical Point Dryer, Quorum, United Kingdom) coated with 8 nm carbon (Q 150 T ES plus, Quorum, United Kingdom) and examined with a scanning electron microscope (EVO 15, Everhart-Thormley-Detector, Zeiss, Oberkochen, Germany) at 8 kV and a sample current of 150 pA.

### Proteome analyses

Bacterial cultures were grown at 30 °C in LB medium for 4 h. Cells were harvested (13 000 x g, 5 min, 4 °C) and resuspended in PBS. Cell pellets were lysed in 60 µl trifluoroacetic acid at 70 °C for 3 minutes, before 600 µl of 2 M Tris Base was added. The cell lysates were cleared by centrifugation for 5 minutes 16 000 x g at 4 °C. Protein concentrations of the cleared lysates were determined by NanoDrop, and 10 µg of the proteins was subjected to tryptic digestion (1:100 for 18 h at 37 °C). The peptides were desalted on Evotip™ using the standard producer protocol and washed twice with 100 µl solvent A (0.1 % TFA (v/v) in water). Peptides were eluted twice in solvent B (80 % ACN, 0.1 % TFA (v/v) in water) and vacuum dried. The samples were resuspended in 20 μL 0.1 % (v/v) formic acid, 1.6 % (v/v) acetonitrile. For each sample 0.4 μL injections were performed. The peptides were separated on a 50 cm μPAC Neo HPLC Column (Thermo Scientific) operating at 50 °C column temperature. Mobile phase A consisted of 0.1 % (v/v) formic acid and mobile phase B of 80 % v/v acetonitrile with 0.1% v/v formic acid. After loading on the column peptides were separated first at a flow rate of 750 nL min^−1^ for 4.95 minutes then 500 nL min^−1^ for 3.15 minutes. Separation was achieved using a linear gradient from 4 % to 8 % mobile phase B over 0.25 minutes, then to 20% over 4.7 minutes, and subsequently to 35 % over 2.25 minutes. The gradient was followed by a rapid increase to 99 % mobile phase B within 0.9 minutes. Eluted peptides were ionized by an EASY-Spray source (Thermo Scientific) and introduced directly into the mass spectrometer. The MS data were acquired in the data-independent (DIA) mode. For precursors with m/z range of 380-980, isolated with 3 m/z isolation windows (0 overlap), resulting in 200 scan events per acquisition cycle. Isolated precursor ions were fragmented using higher-energy collisional dissociation (HCD) with a normalized collision energy of 25 %.MS2 spectra were recorded in the Astral mass analyzer over an m/z range of 150–2000. In parallel, one MS1 spectrum was acquired every 0.6 minutes in the Orbitrap at a resolution of 240 000 (at m/z 200) over an m/z range of 380–980.

### Processing of mass spectrometry data

Raw files obtained from DIA runs were processed using DIA-NN 2.2.0. A FASTA file (UniProt *V. cholerae* serotype O1 reference proteome (Taxon ID: 243277) or *E. coli* (strain K12) reference proteome (Taxon ID: 83333)) was provided. A library-free search was enabled with Trypsin/P protease, 2 missed cleavages, peptide length range 7–30, precursor charge range 1–4, precursor m/z range 300–1800, fragment ion m/z range 200–1800. Cysteine carbamidomethylation was set as a fixed modification. Precursor FDR was set to 1 %, and MBR was activated. All other settings were left as default. Using the resulting “report.tsv” file as input, the data were uploaded to DIA Analyst v0.10.4 (https://analyst-suites.org/apps/dia-analyst/) with 0.05 as an adjusted *p*-value cutoff and 1 as log2 fold change cutoff. Imputation type was set to Perseus type with no normalization. Type of FDR correction was set to Benjamini–Hochberg. The resulting log_2_ fold changes and adjusted p-values were visualized in volcano-style scatter plots generated in Python (pandas, NumPy, matplotlib, seaborn). For each comparison, the log_2_ fold change (sign inverted for plotting) was plotted against the−log_10_-adjusted p-value. Proteins exceeding the fold-change and significance thresholds were highlighted by color. Features with −log_10_-adjusted *p*-values > 4, or alternatively the five most significant hits per regulation direction, were annotated using gene names with collision-avoiding label placement.

## ACKNOWLEDGEMENTS

This work was supported by the University of Hohenheim. We are grateful to the members of the Commichau, Gibhardt and Steuber laboratories for helpful discussions. We thank Luise Weber for the help with some experiments. Support by grants 311211092 (to J.S. and G.F.) and 548311592 (to F.M.C.) from the Deutsche Forschungsgemeinschaft is gratefully acknowledged. S.H. thanks the Landesgraduiertenförderung Baden-Württemberg for financial support. Research reported in this publication was supported by National Institute of Allergy and Infectious Diseases of the National Institutes of Health under award number R03AI180583 (to C.C.H.). We are grateful to Sean Colloms, Daniel Charlier and Tadeusz Kaczorowski for providing plasmids, bacterial strains and helpful advice. We acknowledge the support provided by Susanne Reiße from the Imaging Module of the Core Facility Hohenheim (University of Hohenheim, Stuttgart, Germany) in electron microscopy analysis. Parts of the equipment used was supported by the EFRE EU fund (grant no. 2172959).

## SUPPLEMENTAL MATERIAL

**Supplemental Figure 1.**
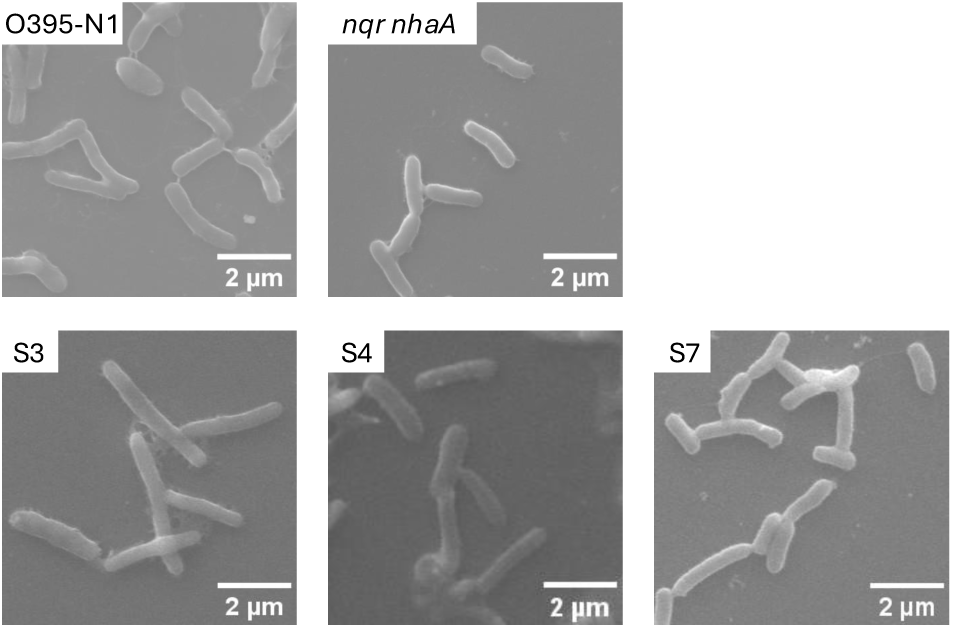
Morphological characterization of *V. cholerae* strains. The strains were grown on LB (pH 6.2, 171 mM NaCl) to exponential phase, harvested and prepared for scanning electron microscopy. **Top panels:** *V. cholerae* O395-N1 (reference strain) and its variant *V. cholerae nqr nhaA* lacking the Na^+^-NQR and the antiporter NhaA. **Bottom panels:** Suppressor mutants S3, S4 and S7 obtained after prolonged cultivation of *V. cholerae nqr nhaA* on LB_Tris_-NaCl (pH 7.8, 150 mM NaCl). All strains were rod shaped and of similar size.

**Supplemental Figure 2.**
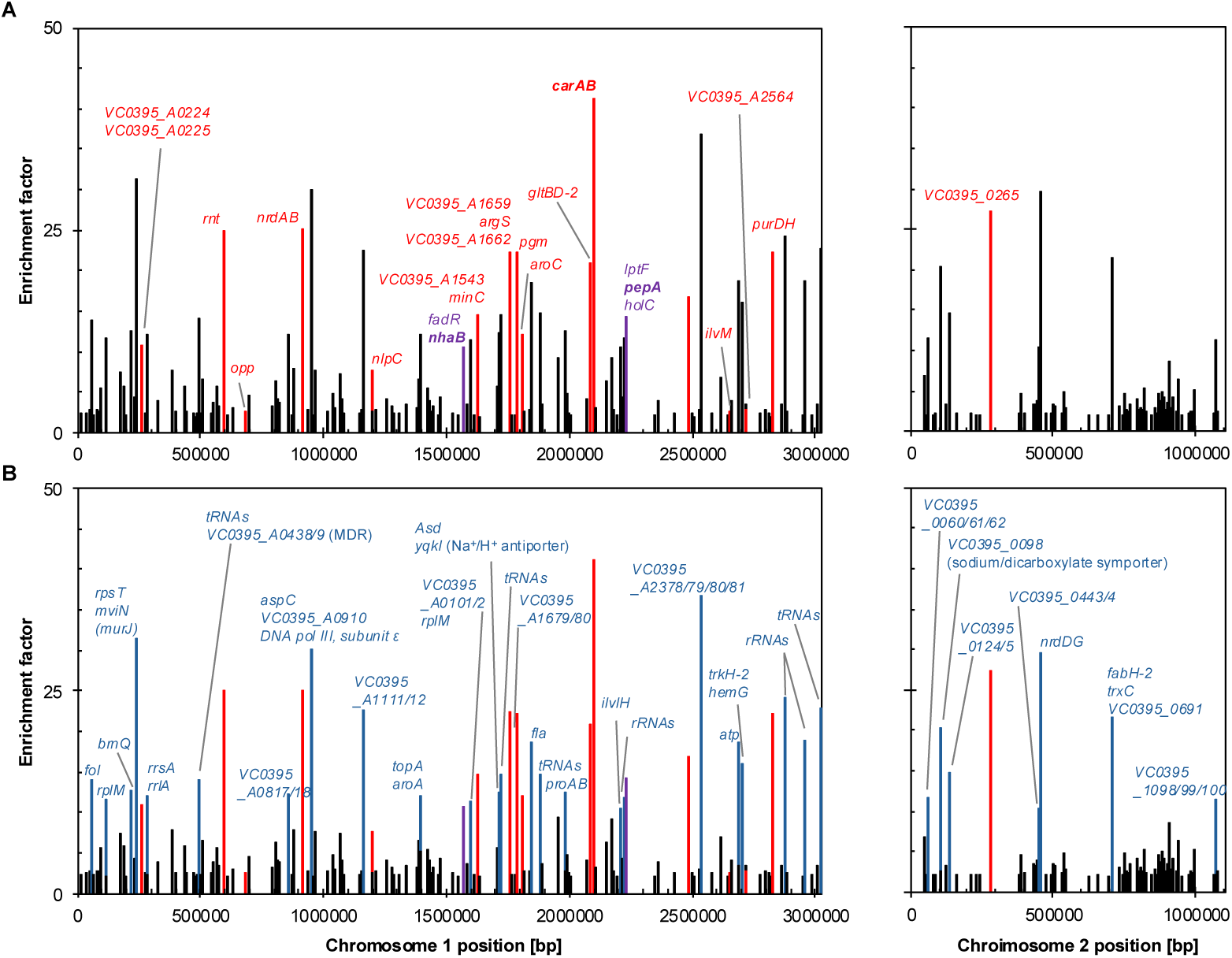
Overlap between PepA binding sites and changes in protein abundance. **(A)** The PepA binding sites in *V. cholerae* O1 biovar El Tor str. N16961 were previously identified by chromatin immunoprecipitation sequencing (ChIP-Seq) [38]. To visualize their distribution across the chromosomes, the PepA binding sites were extracted and mapped onto the the genome of *V. cholerae* O395-N1. The enrichment factors of the binding sites obtained by ChIP-Seq are shown as bars. Red labelled bars indicate PepA binding sites of genes encoding proteins whose abundance was changed only in S3 or S4, or in both suppressors (fold change > 1 or < - 1) (p < 0.05). Purple labelled bars indicate PepA binding sites in *fadR*-*nhaB* and *lptF*-*pepA* intergenic regions, and in the *holC* gene that is located downstream of *pepA*. (**B**) Blue bars indicate PepA binding sites of genes encoding proteins whose abundance remained unchanged.

**Table S1.** Results of proteome analyses.

